# A clinically and genomically annotated nerve sheath tumor biospecimen repository

**DOI:** 10.1101/2019.12.19.871897

**Authors:** Kai Pollard, Jineta Banerjee, Xengie Doan, Jiawan Wang, Xindi Guo, Robert Allaway, Shannon Langmead, Bronwyn Slobogean, Christian F. Meyer, David M. Loeb, Carol D. Morris, Allan J. Belzberg, Jaishri O. Blakeley, Fausto J. Rodriguez, Justin Guinney, Sara J.C. Gosline, Christine A. Pratilas

**Affiliations:** Sidney Kimmel Comprehensive Cancer Center and Department of Oncology, Johns Hopkins University School of Medicine; Sage Bionetworks; Department of Neurology, Johns Hopkins University School of Medicine; Albert Einstein College of Medicine; Department of Orthopedic Surgery, Johns Hopkins University School of Medicine; Department of Neurosurgery, Johns Hopkins University School of Medicine; Department of Pathology, Johns Hopkins University School of Medicine

## Abstract

Nerve sheath tumors occur as a heterogeneous group of neoplasms in patients with neurofibromatosis type 1 (NF1). The malignant form represents the most common cause of death in people with NF1, and even when benign, these tumors can result in significant disfigurement, neurologic dysfunction, and a range of profound symptoms. Lack of human tissue across the peripheral nerve tumors common in NF1 has been a major limitation in the development of new therapies. To address this unmet need, we have created an annotated collection of patient tumor samples, patient-derived cell lines, and patient-derived xenografts, and carried out high-throughput genomic and transcriptomic characterization to serve as a resource for further biologic and preclinical therapeutic studies. In this work, we release genomic and transcriptomic datasets comprised of 55 tumor samples derived from 23 individuals, complete with clinical annotation. All data are publicly available through the NF Data Portal and at http://synapse.org/jhubiobank.

## Background & Summary

Neurofibromatosis type 1 (NF1) is a common neuro-genetic condition caused by mutations in the *NF1* gene. It is characterized by a predisposition to the development of nerve sheath tumors, including cutaneous neurofibromas (cNF), plexiform neurofibromas (pNF), and malignant peripheral nerve sheath tumors (MPNST). Up to 50% of patients with NF1 develop pNF, and 55% of pNF in childhood are symptomatic, resulting in either pain, nerve or organ dysfunction or disfigurement [1]. Currently, surgery is the only treatment option for patients with NF1 who have symptomatic pNF. Progress in the development of nonsurgical therapy for pNF has been limited by a number of factors including: the lack of pNF specific cell culture-based [2, 3] and animal models [4], and limited access to primary tissue from patients with NF1. Although progress is being made in the development and utilization of animal models and cell culture models [5], the limited availability of primary patient tissue remains unaddressed.

To address this gap, we established a local biospecimen repository for the purpose of 1) banking blood fractions and tumor tissue from patients with NF1 undergoing surgical resection of cNF, pNF and/or MPNST at Johns Hopkins Hospital; 2) generating xenograft and cell line models to propagate primary human tissue and cells 3) creating the required infrastructure that supports the sharing of data and tissue resources with the scientific community. The biospecimen repository includes tissue, buffy coat, plasma and serum from patients with NF1 who are undergoing surgical removal of a lesion including, but not limited to, a cNF, diffuse superficial infiltrating neurofibromas, pNF, atypical neurofibromatous neoplasms of uncertain biologic potential (ANNUBP), and MPNST. Each tissue sample has an associated clinical data set and appropriate consent has been obtained to allow sharing of tissue for NF1 research within and outside of Johns Hopkins.

The Johns Hopkins Comprehensive Neurofibromatosis Center (JHCNC) serves a large volume of people with NF1. In concert with the JHCNC, our lab has successfully banked specimens from these patients when they are undergoing surgery, and created and maintained a fully annotated clinical database. All banking procedures are conducted as outlined by the NCI Best Practices (https://biospecimens.cancer.gov/best-practices/index.asp). Specimens are processed and implanted in mice quickly to minimize ischemia time, accurate identification of specimens is ensured by our practices, and all tumor specimens removed from different anatomical locations or different locations within the same tumor or patient are clearly identified and labeled (**Figure 1**).

**Figure 1.**
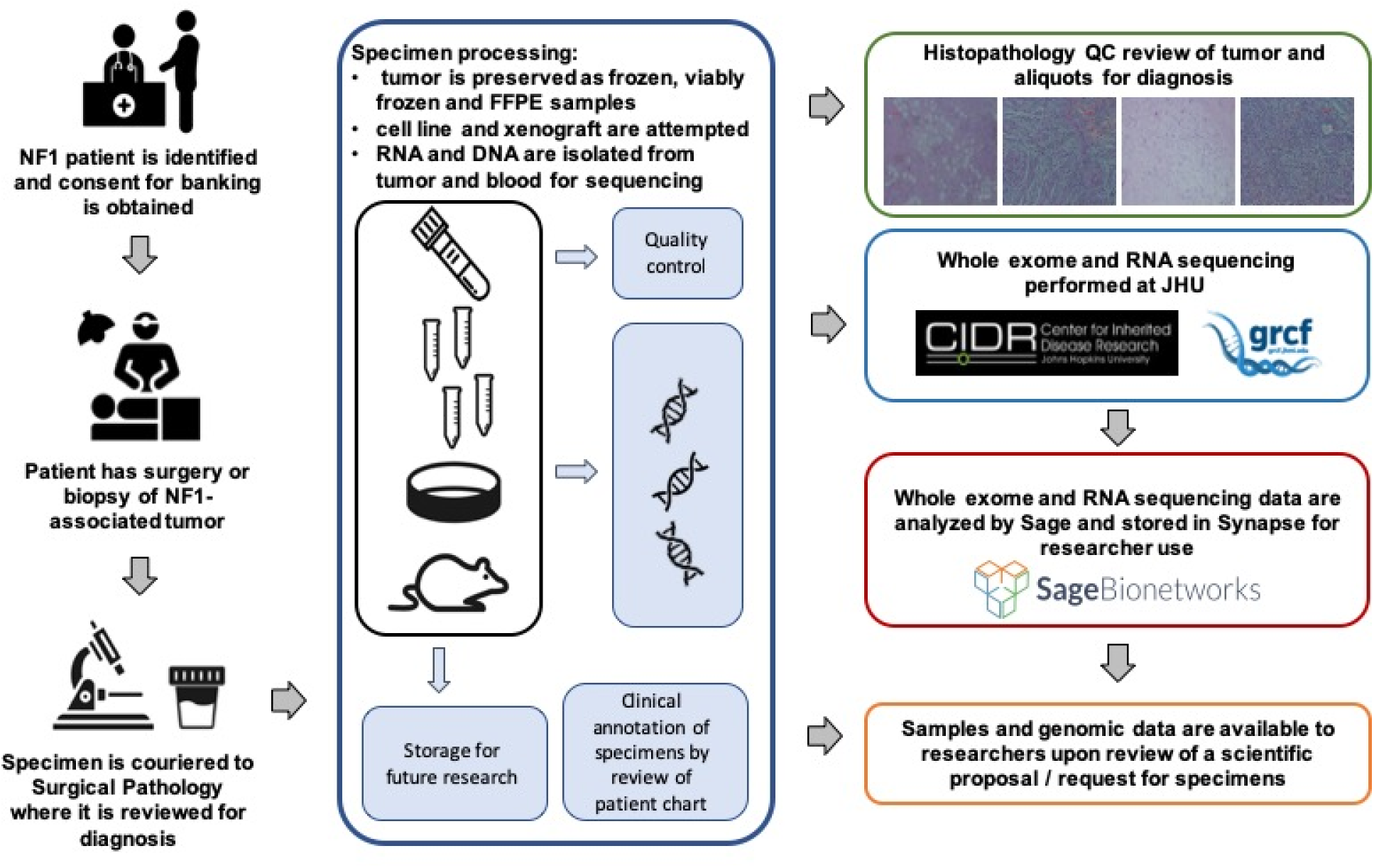
Overview of the Johns Hopkins University NF1 biospecimen repository.

We have leveraged our unique resources, including 1) one of the busiest NF clinics in the country, with specialized physicians and surgeons invested in the process of tissue collection and banking; 2) our expertise in tissue acquisition, handling, and xenograft production, and 3) the comprehensive data-sharing framework of the *NF Data Portal* (http://nfdataportal.org), to establish a key resource and enable sharing of the data generated with the NF1 research community. Currently, we are accruing samples from about twenty patients per year, and expect ongoing accrual at this rate for the foreseeable future.

Herein, we describe the data generated from the tissue bank and shared through the NF Data Portal. We sequenced DNA and/or RNA across 55 tumors from 23 unique patients, as well as any cell lines or xenografts derived from these patient samples. We also sequenced DNA of patient blood cells to include non-tumor (control) genomic data to facilitate downstream analysis. These data are a valuable resource for the NF research community that complements previous NF tumor characterization efforts [6, 7]. All data are available to qualified researchers via the *NF Data Portal* at http://nfdataportal.org to encourage biological exploration and identification of drug targets in NF1.

## Methods

The generation of these data was a close collaboration between the physicians at Johns Hopkins Hospital, the staff of the JH NF1 biospecimen repository, and Sage Bionetworks.

### Patient enrollment

All human subjects research was conducted according to widely accepted practice and under a Johns Hopkins Hospital (JHH) institutional review board (IRB)-approved protocol.

Patients with neurofibromatosis (NF1) having a clinically-indicated surgical resection or biopsy of a NF1-associated tumor (cNF, superficial diffuse infiltrating neurofibroma, pNF, ANNUBP, MPNST) were identified through the review of surgical schedules and communication from the clinical team to the research team at JHH. Patients were reviewed for study eligibility, and then written informed consent was obtained. The JHH IRB-approved consent form includes a description of the voluntary nature of the research and a description of clinical data that will be collected as well as the plan for genetic and genomic analyses and sharing of data. Blood was collected from the majority of patients on the day of surgery.

After successful collection of tumor tissue, the patient’s medical record was reviewed for pertinent demographic information and information related to their NF1 diagnosis (genomic findings, family history, age of diagnosis), characteristics (phenotypic findings, symptoms), and tumor burden (number and size of NF1-associated tumors). These clinical data were stored in a password protected and de-identified database.

### Tumor preservation and quality control

Surgical specimens were couriered to surgical pathology immediately after resection either in saline or in a dry sterile collection cup. The study neuropathologist performed immediate inspection of the tumor to ensure that the sample contained adequate tumor tissue for clinical diagnostic needs. Upon approval, tumor pieces were sampled for banking and transported to the research laboratory in isotonic cell culture medium (RPMI with 20% FBS, supplemented with 1% penicillin-streptomycin and glutamine, PSG).

Specimens were sized into 5-10 mm aliquots under sterile conditions in a biosafety cabinet. Individual aliquots were placed into 10% neutral buffered formalin, cell freezing media (Sigma: C6295), and/or placed into an empty vial and snap frozen on dry ice.

All aliquots were stored with barcoded labels for tracking purposes. Specimens collected from pathologically heterogeneous tumors were embedded in O.C.T. compound (Optimal Cutting Temperature Compound, Fisher Sci 23-730-571) and sectioned into 5 μm sections. One slide from each individual aliquot was reviewed by the study pathologist to confirm histologic diagnosis not only of the tumor as a whole, but that tissue representation and quality was adequate per aliquot.

### Cell culture

A tumor aliquot was mechanically dissociated into a cell suspension in a sterile petri dish filled with supplemented cell culture medium (RPMI, 20% FBS, 1% PSG) using a sterile scalpel. Dissociated cells in culture medium were placed into 75cm^2^ flasks and maintained in an incubator (37°C, 5% CO_2_). Cells were washed and medium changed twice weekly, and passaged until a stable culture was achieved (**Figure 2A-B**). Cells were then viably frozen in Cell Freezing Medium (Sigma C6295) and placed in a Cool Cell temperature controlled freezing chamber at −80°C before transfer to liquid nitrogen for long-term storage.

**Figure 2.**
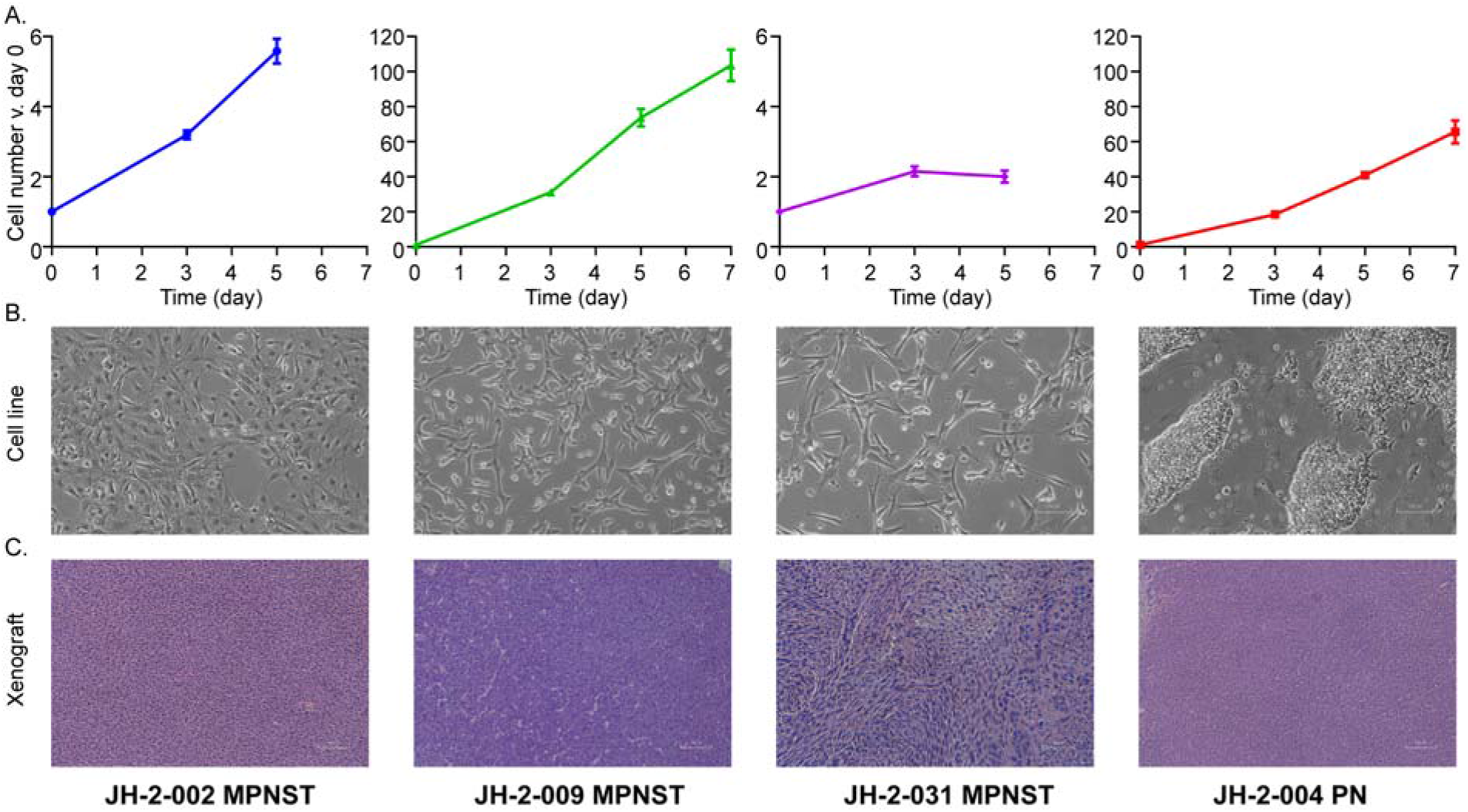
Characterization of patient-derived models. **a.** and **b.** Cells were cultured under standard conditions until the emergence of a consistently replicating population. **a.** Measurement of cell growth rate and calculation of doubling time, shown as a percentage increase over day 0. **b.** 10x photomicrograph of cultured cells at logarithmic growth phase. **c.** Tumor fragments from freshly acquired specimens were implanted subcutaneously into mice; mice were monitored until the development of a palpable tumor. H&E images from four representative patient-derived xenografts (PDX), including three MPNST and one plexiform neurofibroma.

### Patient-Derived Xenograft (PDX) Development

Tissue for PDX generation was maintained in supplemented media (RPMI 20% FBS 1% PSG) until implantation. Adult NOD scid gamma mice (Jackson laboratory: NOD.Cg-Prkdc^scid^II2rg^tm1Wjl^/SzJ (005557)) were anesthetized using a mixture of ketamine and xylazine. Tumor pieces were cut into 2-3 mm fragments and dipped into ice-cold Matrigel (Corning: CB-40230) and immediately implanted into the mouse flank, pre-tibial space, or pre-sacral space.

Two to ten mice were implanted for each tumor where the diagnosis is known to be pNF or MPNST at the time of resection. The overall rate of successful generation of PDX from attempted tumors in this study was 50% for MPNST. Once the tumor reached approximately 15 mm in diameter, tumor was viably frozen in Cell Freezing Medium (Sigma C6295) and a small fragment was passaged into another mouse. Representative H&E images for novel PDX models are shown in **Figure 2C**.

### Exome Sequencing

DNA was isolated from flash-frozen tumor sample using the QIAmp DNA Mini Kit (Qiagen 51304) and quantified and quality confirmed using the NanoDrop 2000 spectrophotometer. Germline DNA was isolated from patient blood, or in the rare case that sufficient blood could not be collected, normal tissue adjacent to tumor was used for germline. Specimens were sequenced by the Genetic Resources Core Facility at Johns Hopkins (https://grcf.jhmi.edu/) and processed similarly to previously published pNF resources [7].

Quality control was carried out via 2% gels, OD_260_ readings, and volume checks upon sample receipt at CIDR (https://www.cidr.jhmi.edu/) to confirm adequate quantity and quality of genomic DNA. Samples were then processed with an Illumina Infinium QCArray-24v1-0 array to confirm sex, identify unexpected duplicates and relatedness, confirm study duplicates and relatedness, provide sample performance information and sample identity confirmation against the sequencing data.

Exome capture was carried out using the Agilent SureSelectXT HumanAllExon V6 (Agilent S07604514) kit. 1μg of genomic DNA was sheared using the Covaris E220 instrument (Covaris) with a shear time of 80 seconds in order to obtain larger insert sizes. A hybrid protocol for library preparation and whole exome enrichment was developed at CIDR (unpublished) based on methods and parameters from Fisher et al., [8] applied to the reagents, volumes and parameters from the Agilent SureSelect XT kit and automated protocol (p/n G7550-90000 revision B). All processing was done in 96 well plate formats using robotics (Beckman FXp, Perkin Elmer Janus, Agilent Bravo, Beckman NX). ‘With Bead’ clean-ups were used following shearing, end repair, A-tailing and adapter ligation. The initial input of GE Healthcare Sera-Mag Magnetic SpeedBeads (Carboxylate-Modified) was based on volumes from the Agilent protocol. After the first clean up, the sample was eluted and the beads remained in the reactions through the final ligation clean-up. These reactions were carried out using the XT reagents, volumes and conditions described in the Agilent protocol. At pre-capture PCR the entire product was amplified, adjusting the water in the reaction to accommodate the increase in DNA sample volume. The PCR enzyme used in all steps was switched from Herculase to HiFi HotStart Ready Mix (Kapa Biosystems) to increase the coverage in GC rich regions. All other aspects follow the Agilent protocol except the number of PCR cycles was increased from 6-8 cycles. 750ng of amplified library was used in an enrichment reaction following Agilent 24-hour hybridization protocol. Post-capture washing was done using the Agilent protocol except the ‘off-bead’ catch process from Fisher et al[8], with a slight change in which samples are not eluted off the DynaBeads (Invitrogen), instead post-capture PCR master mix and indexes are added directly to the beads. Post-capture PCR was done according to the Agilent protocol, with the adjustment of water volume and PCR cycles where needed.

Libraries were sequenced on the HiSeq2500 platform with template generation on the cBot. Conditions included 72 samples per flowcell, 125 base-pair paired-end runs and sequencing chemistry kits HiSeq PE Cluster Kit v4 and HiSeq SBS kit v4.

### Exome-seq variant calling

Single nucleotide variant calling was carried out via a well-tested pipeline with the tools and parameters described in **Table 1**. Copy number alterations were analyzed using the GATK pipeline [9] and additional single nucleotide variant analyses were done using the *DeepVariant* tool (release 0.8) [10].

**Table 1:** List of tools used in exome-seq analysis pipeline

| Analysis Step | Method (version) | Parameters |
| --- | --- | --- |
| Intensity analysis and base calling | Illumina Real Time Analysis (RTA) software (version 1.18.66.4). |  |
| Base call files demultiplexed from a binary format (BCL) to single sample fastq files | CIDRSeqSuite (v 7.5.0, unpublished) |  |
| Fastq file alignment | BWA mem (0.7.15) [20] | 1000 genomes phase 2 (GRCh37) human genome reference |
| Duplicate flagging | Picard (v 2.17.0) |  |
| Base call quality score recalibration and binning (2,10,20,30) | Genome Analysis Toolkit (GATK, v4.0.1.1) |  |
| CRAM file generation | SAMTools (v1.5). [21],<br>GATK (v3.7) [22] | --emitRefConfidence GVCF<br>--max_alternate_alleles 3 |
| SNV Variant Filtering | Variant Quality Score Recalibration (VQSR) method [23] | Annotations of MQRankSum, QD, FS, ReadPosRankSum, MQ and SOR in adaptive error model.<br>HapMap3.3, Omni2.5 and 1000G phase high confidence snp calls used as training sites with HapMap3.3 and Omni2.5 as truth set.<br>0.5% false negative rate |
| Indel Variant Filtering | Variant Quality Score Recalibration (VQSR) method [23] | Annotations of FS, ReadPosRankSum, MQRankSum, QD and SOR in adaptive error model (4 max Gaussians allowed).<br>Curated indels:<br>Mills_and_1000G_gold_standard.indels.b37.vcf<br>1% false negative rate |
| Additional VCF file creation | CalculateGenotypePosteriors | ALL.wgs.phase3_shapeit2_mvncall_integrated_calls.vcf<br>ExAC.r0.3.-sites.vep.vcf 5.20130502.sites.vcf |
| Variant annotation | Annovar (v 2013_02_2) [24] |  |

### RNA-Sequencing

RNA was isolated from flash-frozen tumor specimens by grinding each with a mortar and pestle while frozen with liquid nitrogen. RNA was isolated using the RNeasy Mini Kit (Qiagen Cat No: 74104) and quantified and quality confirmed via a NanoDrop 2000 spectrophotometer. For each sample, paired-end RNA-Seq data run in eight lanes was concatenated into two fastq files. For each sample, paired-end RNA-Seq data were converted from bam files to two fastq files using *BEDtools* [11]. Quality control was performed using *FastQC* and combined into one file utilizing *MultiQC* [12]. The report was uploaded to Synapse (syn17095976). One sample (2-025 Neurofibroma) was found to have high a percentage of duplication (> 90%) and was therefore removed from the study, which leaves 28 samples in total. Alignment was performed using *Salmon* (0.11.3) [13] and aligned with *gencode* (version 29). The raw counts matrix was assembled by importing the output files of Salmon alignment via *tximport*. The alignment output files and the raw counts matrix were uploaded to Synapse (syn19522967). A harmonized version of the RNA-seq dataset for 28 samples across 22 individuals is available at syn20812185 and is also harmonized with previously-sequenced pNF cell culture data [7]. The analysis code is freely available at https://github.com/Sage-Bionetworks/JHU-biobank.

## Code availability

A Github repository (http://github.com/sage-bionetworks/JHU-biobank) contains the codes required to generate the figures. The tutorials are provided in R and Python languages, contained in the `r_demos` and `py_demos` directories respectively. All of the analytical code is provided in the directory marked “`analysis`”. Additionally, we have provided Docker containers and R scripts to facilitate reproducibility of the figures in the paper.

## Data Records

All de-identified data can be retrieved from the Synapse project site at http://synapse.org/jhubiobank. The project page describes the details of the project as well as how to gain access to the data. At the time of this publication, the site includes RNA and DNA data from 23 patients, as described in **Table 2**. Clinical metadata is available on the Synapse data site at http://synapse.org/jhubiobank. Samples characterized via exome-seq are described in **Table 3**. Samples characterized by RNA-seq are described in **Table 4**.

**Table 2:**
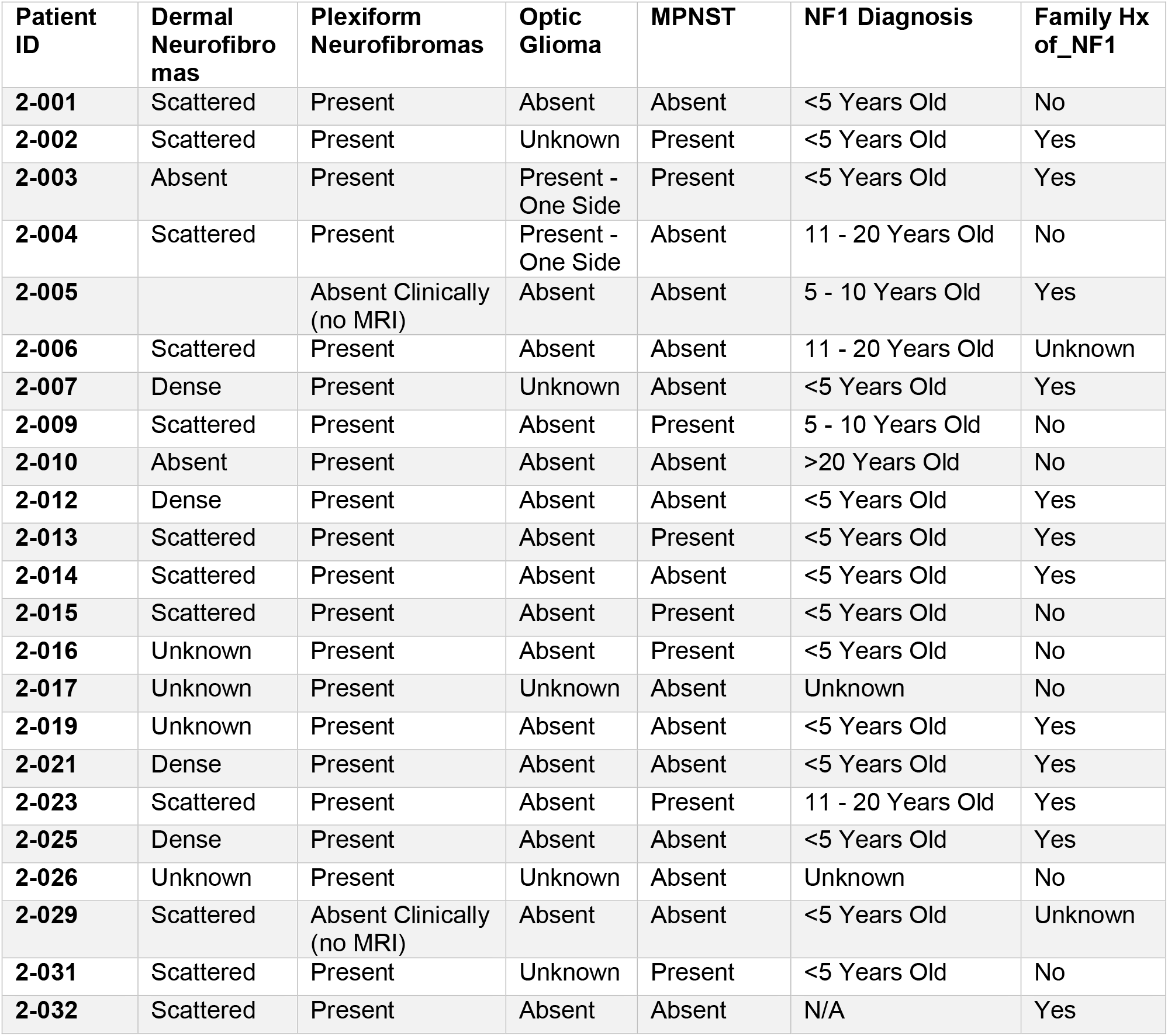
List of patients and clinical variables

**Table 3:** Samples characterized via exome-seq

| Patient ID | sex | source | Cell Line | PDX | specimenID |
| --- | --- | --- | --- | --- | --- |
| 2-001 | male | Blood |  |  | 2-001 Blood |
| 2-001 | female | Plexiform Neurofibroma |  |  | 2-001 Plexiform Neurofibroma |
| 2-002 | male | Blood |  |  | 2-002 Blood |
| 2-004 | female | Blood |  |  | 2-004 Blood |
| 2-004 | female | Neurofibroma | X |  | 2-004 NF Cell Line |
| 2-004 | female | Plexiform Neurofibroma |  |  | 2-004 Plexiform Neurofibroma |
| 2-004 | female | Plexiform Neurofibroma | X |  | 2-004 pNF Cell Line |
| 2-005 | female | Blood |  |  | 2-005 Blood |
| 2-005 | female | Neurofibroma |  |  | 2-005 Neurofibroma |
| 2-006 | female | Blood |  |  | 2-006 Blood |
| 2-006 | female | Plexiform Neurofibroma |  |  | 2-006 Plexiform Neurofibroma |
| 2-009 | male | Blood |  |  | 2-009 Blood |
| 2-010 | male | Blood |  |  | 2-010 Blood |
| 2-010 | male | Neurofibroma |  |  | 2-010 Neurofibroma |
| 2-012 | male | Blood |  |  | 2-012 Blood |
| 2-012 | male | Neurofibroma |  |  | 2-012 Neurofibroma |
| 2-013 | female | Blood |  |  | 2-013 Blood |
| 2-013 | female | Plexiform Neurofibroma |  |  | 2-013 Plexiform Neurofibroma |
| 2-014 | male | Blood |  |  | 2-014 Blood |
| 2-014 | male | Neurofibroma |  |  | 2-014 Neurofibroma |
| 2-015 | male | Blood |  |  | 2-015 Blood |
| 2-015 | male | Neurofibroma |  |  | 2-015 Neurofibroma |
| 2-015 | male | Plexiform Neurofibroma |  |  | 2-015 Plexiform Neurofibroma |
| 2-016 | female | Blood |  |  | 2-016 Blood |
| 2-016 | female | Neurofibroma |  |  | 2-016 Neurofibroma |
| 2-017 | male | Blood |  |  | 2-017 Blood |
| 2-017 | male | Neurofibroma |  |  | 2-017 Neurofibroma |
| 2-019 | female | Blood |  |  | 2-019 Blood |
| 2-019 | female | Neurofibroma |  |  | 2-019 Neurofibroma |
| 2-021 | male | Blood |  |  | 2-021 Blood |
| 2-021 | male | Neurofibroma |  |  | 2-021 Neurofibroma |
| 2-023 | male | Blood |  |  | 2-023 Blood |
| 2-023 | male | Plexiform Neurofibroma |  |  | 2-023 Plexiform Neurofibroma |
| 2-025 | female | Blood |  |  | 2-025 Blood |
| 2-025 | female | Neurofibroma |  |  | 2-025 Neurofibroma |
| 2-026 | female | Blood |  |  | 2-026 Blood |
| 2-026 | female | Neurofibroma |  |  | 2-026 Neurofibroma |
| 2-029 | male | Blood |  |  | 2-029 Blood |
| **2-029** | male | Neurofibroma |  |  | 2-029 Neurofibroma |
| **2-031** | male | Blood |  |  | 2-031 Blood |
| **2-031** | male | Malignant Peripheral Nerve Sheath Tumor | X |  | 2-031 Cell Line |
| **2-031** | male | Malignant Peripheral Nerve Sheath Tumor |  |  | 2-031 Malignant Peripheral Nerve Sheath Tumor |
| **2-031** | male | Malignant Peripheral Nerve Sheath Tumor |  | X | 2-031 Xenograft |
| **2-032** | female | Plexiform Neurofibroma |  |  | 2-032 Plexiform Neurofibroma |
| **2-032** | female | Skin |  |  | 2-032 Skin |

**Table 4:** Samples characterized via RNA-seq

| **Patient ID** | **sex** | **Source** | **Cell Line** | **PDX** | **specimenID** |
| --- | --- | --- | --- | --- | --- |
| **2-001** | female | Plexiform Neurofibroma |  |  | 2-001 Plexiform Neurofibroma |
| **2-002** | male | Malignant Peripheral Nerve Sheath Tumor | X |  | 2-002 Cell Line |
| **2-002** | male | Malignant Peripheral Nerve Sheath Tumor |  |  | 2-002 Malignant Peripheral Nerve Sheath Tumor |
| **2-002** | male | Malignant Peripheral Nerve Sheath Tumor |  | X | 2-002 xenograft |
| **2-003** | male | Plexiform Neurofibroma |  |  | 2-003 Plexiform Neurofibroma |
| **2-004** | female | Plexiform Neurofibroma |  |  | 2-004 Plexiform Neurofibroma |
| **2-004** | female | Plexiform Neurofibroma | X |  | 2-004 pNF Cell Line |
| **2-004** | female | Plexiform Neurofibroma |  | X | 2-004 xenograft |
| **2-005** | female | Neurofibroma |  |  | 2-005 Neurofibroma |
| **2-006** | female | Blood |  |  | 2-006 Blood |
| **2-006** | female | Plexiform Neurofibroma |  |  | 2-006 Plexiform Neurofibroma |
| **2-007** | male | Neurofibroma |  |  | 2-007 Neurofibroma |
| **2-009** | male | Malignant Peripheral Nerve Sheath Tumor | X |  | 2-009 Cell Line |
| **2-009** | male | Malignant Peripheral Nerve Sheath Tumor |  |  | 2-009 Malignant Peripheral Nerve Sheath Tumor |
| **2-009** | male | Malignant Peripheral Nerve Sheath Tumor |  | X | 2-009 Xenograft |
| **2-010** | male | Neurofibroma |  |  | 2-010 Neurofibroma |
| **2-012** | male | Blood |  |  | 2-012 Blood |
| **2-012** | male | Neurofibroma |  |  | 2-012 Neurofibroma |
| **2-013** | female | Blood |  |  | 2-013 Blood |
| **2-013** | female | Plexiform Neurofibroma |  |  | 2-013 Plexiform Neurofibroma |
| **2-014** | male | Blood |  |  | 2-014 Blood |
| **2-014** | male | Neurofibroma |  |  | 2-014 Neurofibroma |
| **2-016** | female | Neurofibroma |  |  | 2-016 Neurofibroma |
| **2-017** | male | Blood |  |  | 2-017 Blood |
| **2-017** | male | Neurofibroma |  |  | 2-017 Neurofibroma |
| **2-019** | female | Blood |  |  | 2-019 Blood |
| **2-019** | female | Neurofibroma |  |  | 2-019 Neurofibroma |
| **2-021** | male | Neurofibroma |  |  | 2-021 Neurofibroma |
| **2-023** | male | Malignant Peripheral Nerve Sheath Tumor |  |  | 2-023 Malignant Peripheral Nerve Sheath Tumor |
| **2-025** | female | Neurofibroma |  |  | 2-025 Neurofibroma |
| **2-026** | female | Blood |  |  | 2-026 Blood |
| **2-026** | female | Neurofibroma |  |  | 2-026 Neurofibroma |
| **2-029** | male | Blood |  |  | 2-029 Blood |
| **2-029** | male | Neurofibroma |  |  | 2-029 Neurofibroma |
| **2-031** | male | Malignant Peripheral Nerve Sheath Tumor |  |  | 2-031 Malignant Peripheral Nerve Sheath Tumor |
| **2-032** | female | Plexiform Neurofibroma |  |  | 2-032 Plexiform Neurofibroma |

## Technical Validation

We evaluated the genetic fidelity of the PDX and cell line models by comparing mutational profiles of commonly mutated genes [14–19] from the exome-seq data to showcase that the mutational profiles between the original tumor were maintained in the cell lines and the PDX models, depicted in **Figure 3a**.

**Figure 3.**
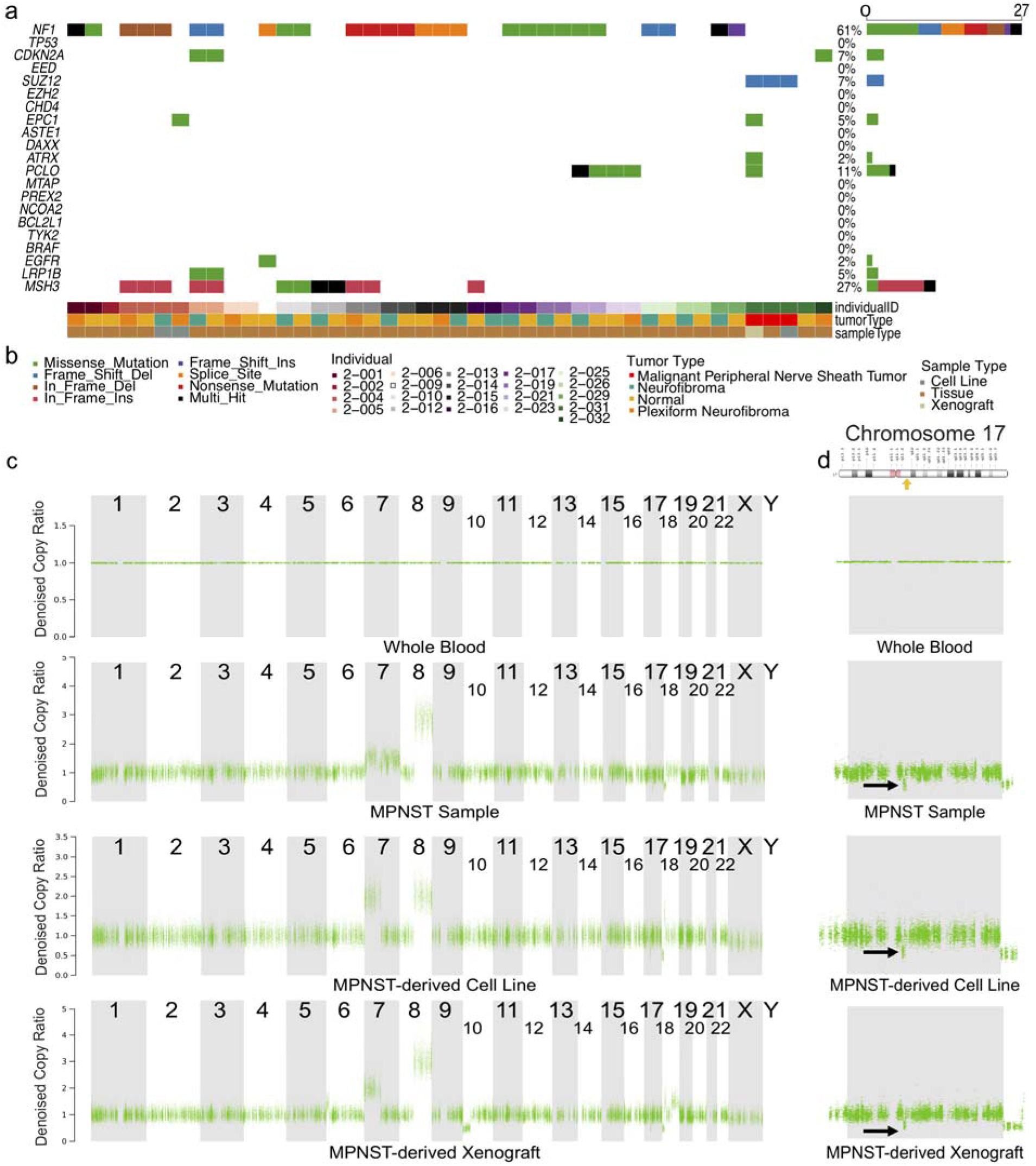
Genomic profile of patient blood, tumor, and patient-derived cell line and xenograft samples. **a.** represents genomic alterations in commonly mutated genes across all samples for which there are sequencing data. Gene names are listed along the left, with the percent of samples in which that gene is mutated on the right. Sample metadata are located at the bottom of the figure. Common variants not included in the plot. **b.** Legend for panel a. **c.** Plots from copy ratio analysis of all chromosomes in the four samples derived from patient 2-031. **d.** Top panel shows a diagrammatic representation of chromosome 17 with NF1 locus highlighted by a yellow arrow (adapted from https://ghr.nlm.nih.gov/gene/NF1#location). The bottom four panels are high resolution visualizations of chromosome 17 in 2-031 specimens showing the presence of a reduction in copy ratio at the NF1 locus (indicated by black arrows) in the MPNST tumor sample, the derived cell line, and the xenograft.

In addition to measuring single nucleotide variants (SNVs), we also measured copy ratio alterations to ensure that the cell line and xenograft models recapitulated the genomic profile of the original sample. **Figure 3c** shows plots of the copy ratios of all chromosomes in the various samples derived from patient 2-031. **Figure 3d** shows a higher resolution plot of the copy ratio specifically in Chromosome 17. There was a detectable reduction in copy ratio (CR) near the NF1 locus (CR of blood = 1, CR of tumor = 0.5). This reduction in copy ratio around the NF1 locus in the tumor tissue, indicated by the black arrow, was preserved across the cell line and xenograft models derived from the same patient.

For technical validation of RNA-seq data, the quality of RNA-seq data was re-evaluated in-silico after the initial quality control during sequencing. The z-scored total counts per gene were plotted (**Figure 4a)** to show that the number of reads per gene were similarly distributed across all samples thus confirming that samples were of analytical quality. The 2-004 xenograft was found to be an outlier sample with generally lower reads than the others. A principal component analysis (PCA) of the samples (shown in **Figure 4b**) confirmed that the cell lines were similar to the xenografts but both showed some drift from the original tissue samples.

**Figure 4.**
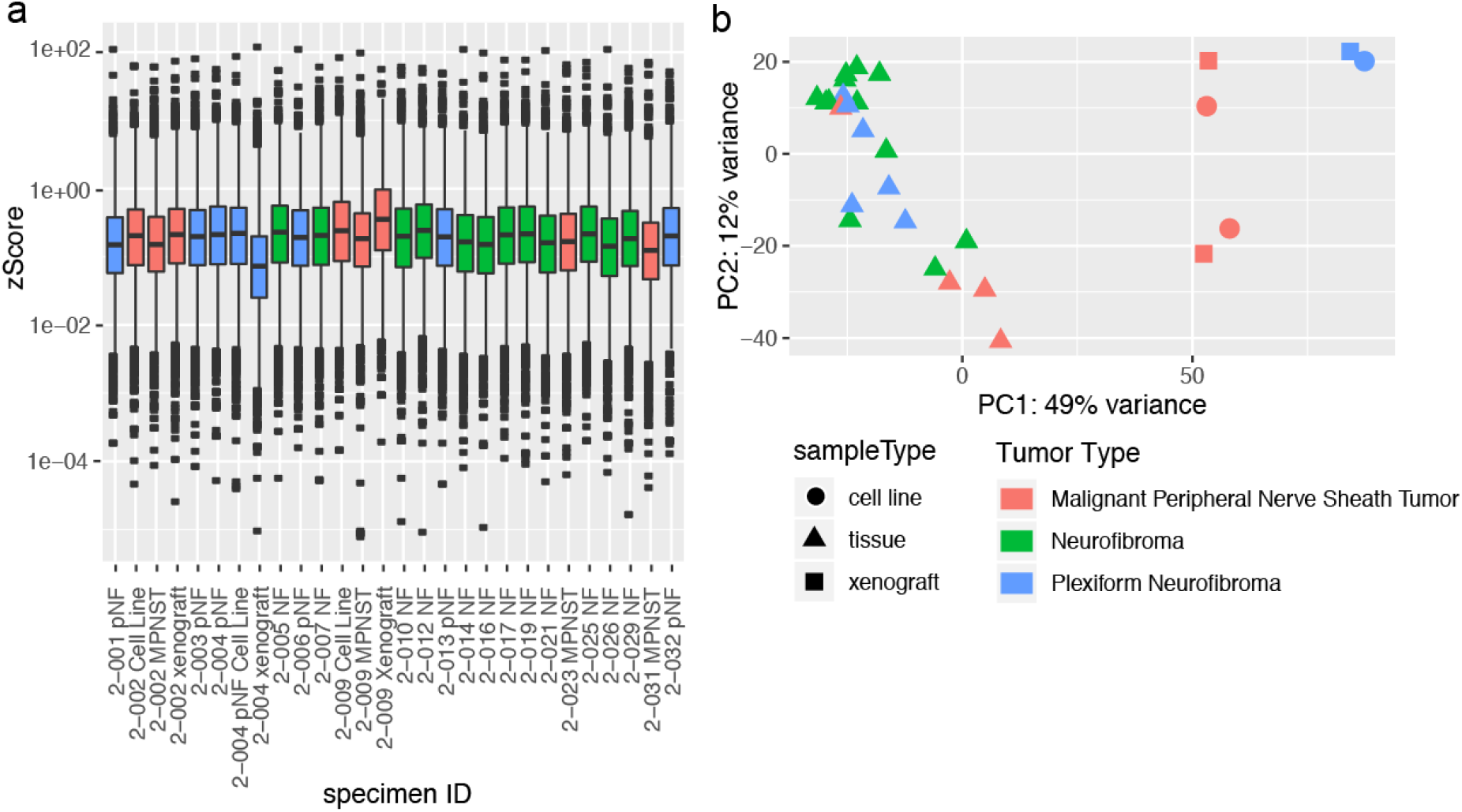
Technical validation of RNA-seq data. **a.** Boxplot of normalized counts (zScores) for each gene for each dataset. **b.** Depicts the first two principal components of each sample, colored by tumor type. Shape represents whether the sample is a cell line (circle), xenograft (square), or tumor tissue (triangle).

## Usage Notes

All data can be found on the Biobank Synapse project page at http://synapse.og/jhubiobank. The data is freely available to qualified researchers upon request for access, following the instructions described on the project page.

## Acknowledgements

The JH NF1 biospecimen repository is supported by a grant from the Neurofibromatosis Therapeutic Acceleration Program (NTAP, http://www.n-tap.org/) to C.A.P. Analysis by Sage Bionetworks is supported through the Neurofibromatosis Therapeutic Acceleration Program (NTAP, http://www.n-tap.org/).

## Author contributions

K.P.: specimen handling and clinical annotation, oversight of data integrity

D.M.L., J.O.B., and C.A.P.: concept, design and scientific oversight

J.O.B., B.L.S., S.M.L. C.F.M., C.D.M. and A.J.B.: patient care and acquisition of specimens

J.O.B., B.L.S., S.M.L. and F.J.R.: assignment of clinical diagnosis, verification of accuracy of specimen diagnosis

X.G, X.D, J.B, R.A. and S.G.: data curation and bioinformatic analysis

K.P., J.B., J.W., C.P., and S. G.: generation of figures

C.A.P. and S.G.: manuscript writing

All authors: final review of manuscript

